# A temperature sensitive mutant screen reveals translational stress-induced cell cycle regulation in a thermophilic archaeon

**DOI:** 10.1101/2025.01.07.631727

**Authors:** Sherman Foo, Yin-Wei Kuo, Jovan Traparić, Dennis W. Grogan, Buzz Baum

**Affiliations:** Medical Research Council Laboratory of Molecular Biology; Cambridge, United Kingdom; University of Cincinnati, Department of Biological Sciences, Ohio, United States of America

**Author notes:** These authors contributed equally to this work (Sherman Foo, Yin-Wei Kuo).

## Abstract

The homology of the archaeal and eukaryotic ribosome provides one of the key pieces of evidence that underpins the idea that eukaryotes acquired their core information processing machinery from archaea. Since this discovery, reverse genetics has been used to study the functions of many archaeal proteins with eukaryotic homologues. Yet, our general understanding of archaeal growth and division remains unclear, in part because of difficulties of carrying out unbiased genetic screens in archaea. Here, by overcoming several technical hurdles we have used a screen of temperature sensitive mutants in *Sulfolobus acidocaldarius* to identify core regulators of cell growth and division. First, flow cytometry was used to define DNA content, identifying a set of mutants defective in cell cycle progression at elevated growth temperatures. Using genome sequencing and plasmid rescue, we then identified a point mutation in the large ribosomal subunit that inhibits translation and prevents entry into division following a shift to the restrictive temperature. This study reveals a link between translation and cell cycle control, and opens up the future possibility of using forward genetic screens in archaea to further our understanding of the similarities and differences in the cell biology of archaea, bacteria and eukaryotes.

**Significance statement:**

- Current knowledge of archaeal cell biology is limited by the lack of forward genetics.
- Whole genome sequencing and plasmid rescue identifies causative mutation in a temperature sensitive mutant strain.
- A mutation in a ribosomal subunit blocks translation to prevent entry into division.

## Introduction

Archaea have cell biological characteristics that are similar and different to both bacteria and eukaryotes. For example, while archaea have a circular DNA like bacteria, their transcription and translation machinery are more akin to those of eukaryotes (Lake *et al*., 1984; Iwabe *et al*., 1989; Cezanne *et al*., 2024), and they have membranes unique to their domain (Boucher *et al*., 2004; Albers and Meyer, 2011). While most recent work in archaea has focused on proteins with homologues in eukaryotes, bacteria or viruses whose functions are well-defined (Iwai *et al*., 2000; Grabowski and Kelman, 2001; Lindås *et al*., 2014; Liu *et al*., 2017; Meyer *et al*., 2022; Zheng *et al*., 2023), such an approach necessarily overlooks the many conserved archaea proteins with unknown function. This leads to significant bias in our view of archaeal cell biology, since archaea were only identified as a distinct domain of life in the 1970s (Woese and Fox, 1977) - long after the discovery of eukaryotes and bacteria. As an example of this, since Sulfolobales lack obvious Cyclin-Dependent Kinases and cyclin homologs, it remains unclear how they organise and control their DNA replication and cell division cycles (Bernander and Poplawski, 1997; Hjort and Bernander, 1999).

*Sulfolobus acidocaldarius*, along with related species from the family of Sulfolobaceae, are members of the TACK archaea, and are currently the closest experimentally tractable relatives of eukaryotes. In recent years, these organisms have emerged as excellent models for studying the cell biology of archaea due to their rapid growth, the expanding set of genetic tools, and their stable genome organization (Grogan *et al*., 2001; Brügger *et al*., 2002). Although there have been few attempts at forward genetics, in 2000, using a classical forward genetics approach, temperature sensitive (TS) mutants of *S. acidocaldarius* were isolated by Bernander, Grogan, Mehdigholi and Poplawski (Bernander *et al*., 2000). This identified strains that exhibited defects in chromosome replication, DNA segregation, growth and cell division when cultured at the non-permissive temperature. However, due to technical challenges, it was not possible at the time to identify the mutations responsible for these phenotypes.

Since then, there have been substantial advances in genomic sequencing and bioinformatics tools. The *S. acidocaldarius* genome has now been sequenced (Chen *et al*., 2005) – along with the genomes of the related *Saccharolobus islandicus* and *Saccharolobus solfataricus* (She *et al*., 2001; Zhang *et al*., 2018). Furthermore, in recent years, developments in molecular biology, genetics (Wagner *et al*., 2012; Zink *et al*., 2019) and imaging techniques (Pulschen *et al*., 2020; Charles-Orszag *et al*., 2021; Cezanne *et al*., 2023) have expanded the toolbox for the study and characterization of archaea. Given this, we thought it is time to revisit the earlier screen in the hope of identifying and characterising the genes regulating key events in the archaeal cell cycle.

## Results

### Identifying cell cycle mutants through temperature sensitive mutant screen

A set of temperature sensitive mutants of *S. acidocaldarius* were previously isolated in a classical forward genetic screen in 2000 (Bernander *et al*., 2000). In this study, chemical mutagenesis of *S. acidocaldarius* was performed utilizing methylnitroninitrosoguanidine. Temperature sensitive mutants were then identified via replica plating at a permissive and restrictive temperature (Fig. 1A). Note that the screen differed from cell cycle screens in yeast, which typically focused on mutants that were not strongly lethal at restrictive temperature (Hoffman *et al*., 2015; Adames *et al*., 2019). Although the forward genetic screen successfully identified a number of mutant lines displaying cell division defects at the restrictive temperature, the causative mutations giving rise to these conditional phenotypes were not identified due to technological limits at the time (Bernander *et al*., 2000). With a more complete set of genetic tools in Sulfolobus and with the advancement in next generation sequencing, we decided to revisit the hits from the screen with the aim of identifying the mutations that perturb cell division cycle in a temperature-sensitive manner.

**Figure 1.**
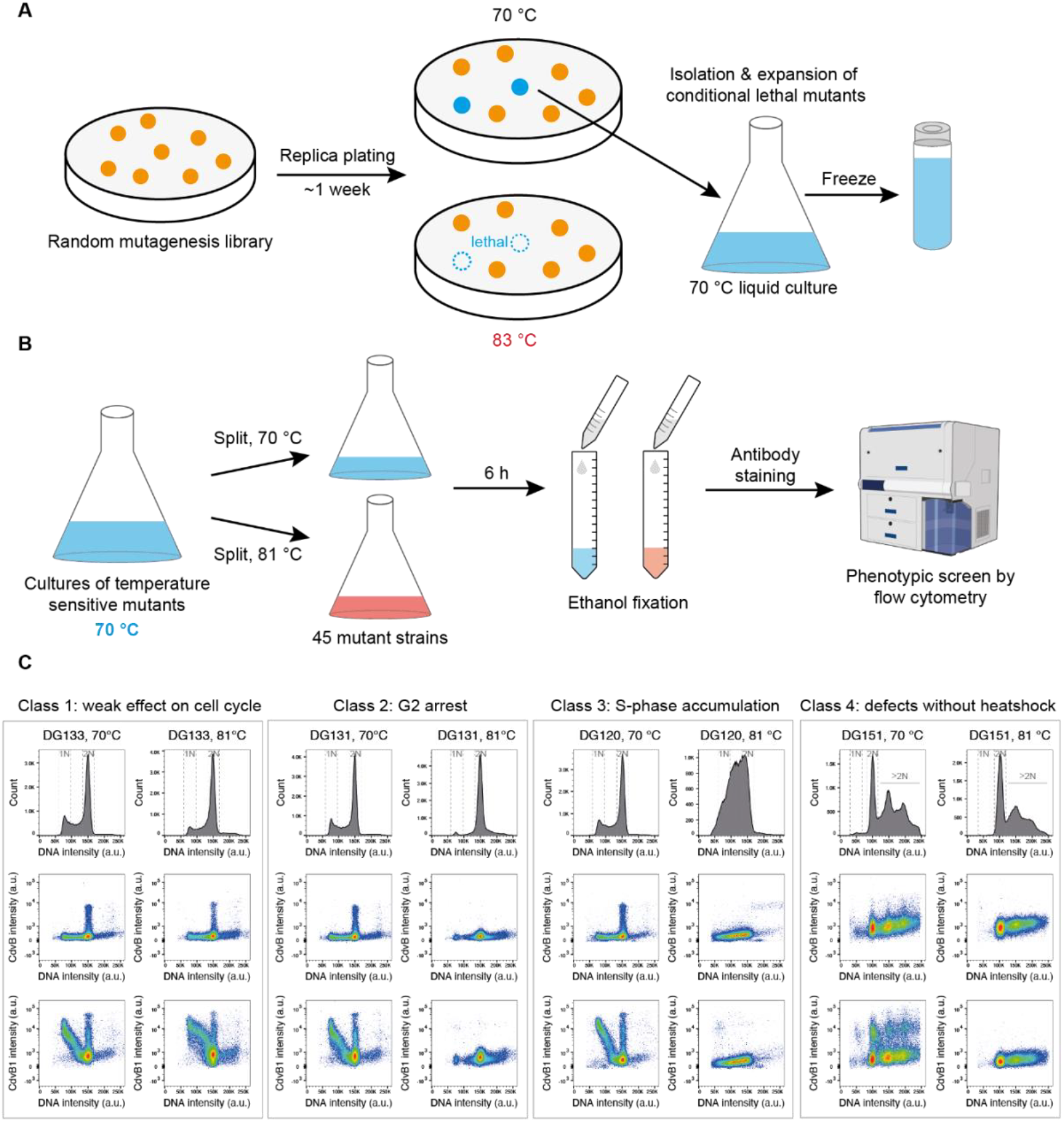
Visualising cell division phenotypes through flow cytometry analysis. **(A)** Schematic showing the temperature sensitive mutant screen performed by Bernander and colleagues (Bernander *et al*., 2000). Chemical mutagenesis of *S. acidocaldarius* was performed, followed by replica plating at the permissive (70 °C) and restrictive temperatures (83 °C). TS mutants were isolated and frozen. **(B)** Rescreening for cell cycle and cell division phenotypes via flow cytometry analysis of the DNA content of asynchronous cultures was performed in this study, following 6 hours of culture at the permissive and lower restrictive temperatures (81 °C). **(C)** Representative graphs showing DNA content and ESCRT-III protein levels of the four classes of cell cycle mutants following 6 hours of culture at the restrictive condition; n=1×10^5^ events.

To re-screen these isolated mutants, we cultured strains at the permissive (70 °C) and restrictive temperatures (81 °C) for 6 hours before ethanol fixation (Fig. 1B). To identify mutants with potential cell cycles or cell division defects, we then analysed the relative populations of cells at different cell cycle stages using flow cytometry to assess DNA content and the levels of ESCRT-III proteins (CdvB, CdvB1). This identified four phenotypic classes of mutant (Fig 1C, and see summary in supplementary table S1). The majority of these (22/44 mutants) had a mild cell cycle phenotype 6 hours after a shift to the restrictive temperature (equivalent to 1 to 2 cell cycles), which was characterised by a modest decrease in the ratio of the cell population in G1 and G2 of the cell cycle (Class 1). A number of mutants (11/44 mutants) suffered a G2 arrest following the temperature shift (Class 2) - a phenotype that was accompanied by a strong reduction in the expression of CdvB and CdvB1. Three mutants displayed an increase in the S-phase population (Class 3). Lastly, two mutant lines were characterised by cells with multiple copies of DNA at both the permissive and restrictive temperature (Class 4). Conditional lethal mutants (6 strains) were also identified that thrived at the permissive temperature of 70 °C, but did not survive 6 hours of culture at the restrictive temperature of 81 °C. A shorter incubation time or a lower restrictive temperature might be needed to assess the impact on their cell cycle process.

### Reversibility of temperature sensitive phenotypes

Our initial re-screen identified several temperature-sensitive mutant strains that showed G2 arrest at the restrictive temperature. We subjected two of these mutants, DG155 and DG131, to further investigation to determine whether the phenotype is reversible. For this experiment, cells arrested at G2 at restrictive temperature (81 °C) were followed after shifting back to permissive temperature (70 °C). A flow cytometric analysis of DNA content showed that both strains displayed an increase in G2 cells after 4.5 hours of incubation at the restrictive temperature of 81 °C (Fig. 2A and 2B). After shifting back to the permissive temperature of 70 °C, only DG155 resumed cycling, leading to the accumulation of a population of G1-S cells (Fig. 2A, black curve in 2C). By contrast, the vast majority of DG131 cells remained stuck in G2 three hours after the shift back to 70 °C (Fig. 2B, magenta curve in 2C).

**Figure 2.**
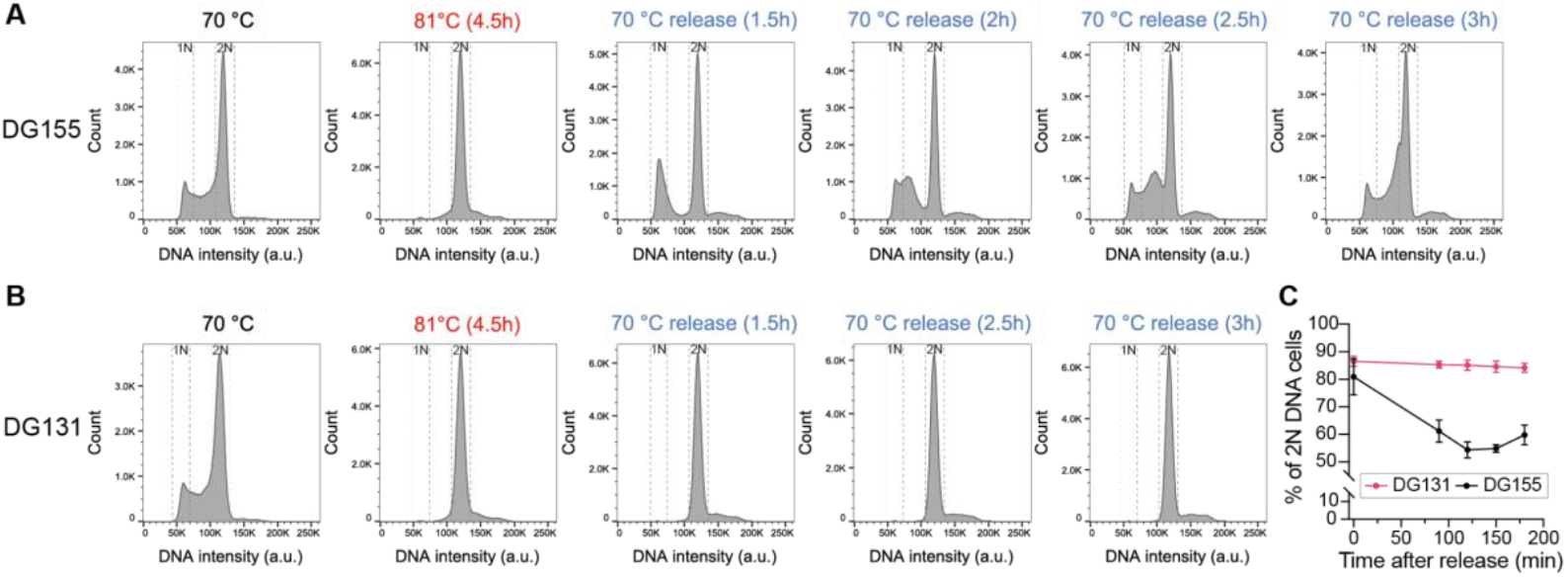
Reversibility of temperature sensitive mutants. DG155 **(A)** and DG131 **(B)** were heated to non-permissive temperature (81 °C) for 4.5 hr and shifted back to permissive temperature (70 °C). Cells at different cell cycle stages were monitored by histogram of DNA intensity using flow cytometry; n=2.5×10^5^ events. Both DG155 and DG131 showed G2 arrest after 4.5 hours incubation at 81 °C, while only DG155 showed obvious release. **(C)** Quantification of cells with 2N DNA content after shifting from 81 °C back to 70 °C. Error bars: SDs; N=3 independent cultures.

### An E34K point mutation in the large ribosomal protein subunit Rpl1 inhibits translation at the restrictive temperature

To identify the mutations responsible for the cell cycle arrest in each case, whole genome sequencing was performed for several selected strains from phenotype classes Class 2, 3 and 4. This identified multiple point mutations per strain when compared to the reference genome in the NCBI database (GCF_002215565.1). The sequencing results are summarised in supplementary table S2.

In order to identify the mutation responsible for the cell cycle arrest in one case, we focused on DG131, a class 2 mutant. This mutant line was chosen for several reasons: (i) the cell cycle arrest phenotype was robust and not sensitive to the heating speed nor the incubation time at the restrictive temperature; (ii) the cells were irreversibly arrested at G2, suggesting the mutation likely has a strong effect; (iii) the number of unique missense mutations present in the genome (i.e., not found in other sequenced TS mutants) were fewer in this strain relative to those seen in other TS strains.

After further inspection of the genome, we identified a point mutation E34K in the large ribosomal protein subunit Rpl1 (Saci1548; Uniprot accession P35024) as a promising candidate for further investigation. The mutation is located at the interface between Rpl1 and 23S rRNA (Fig. 3A) and the E34 side chain of Rpl1 was found to form a hydrogen bond with G2262 on 23S rRNA (Fig. 3B) (Nikulin *et al*., 2003). The E34K mutation reverses the charge, likely weakening the interaction between Rpl1 and the 23S rRNA.

**Figure 3.**
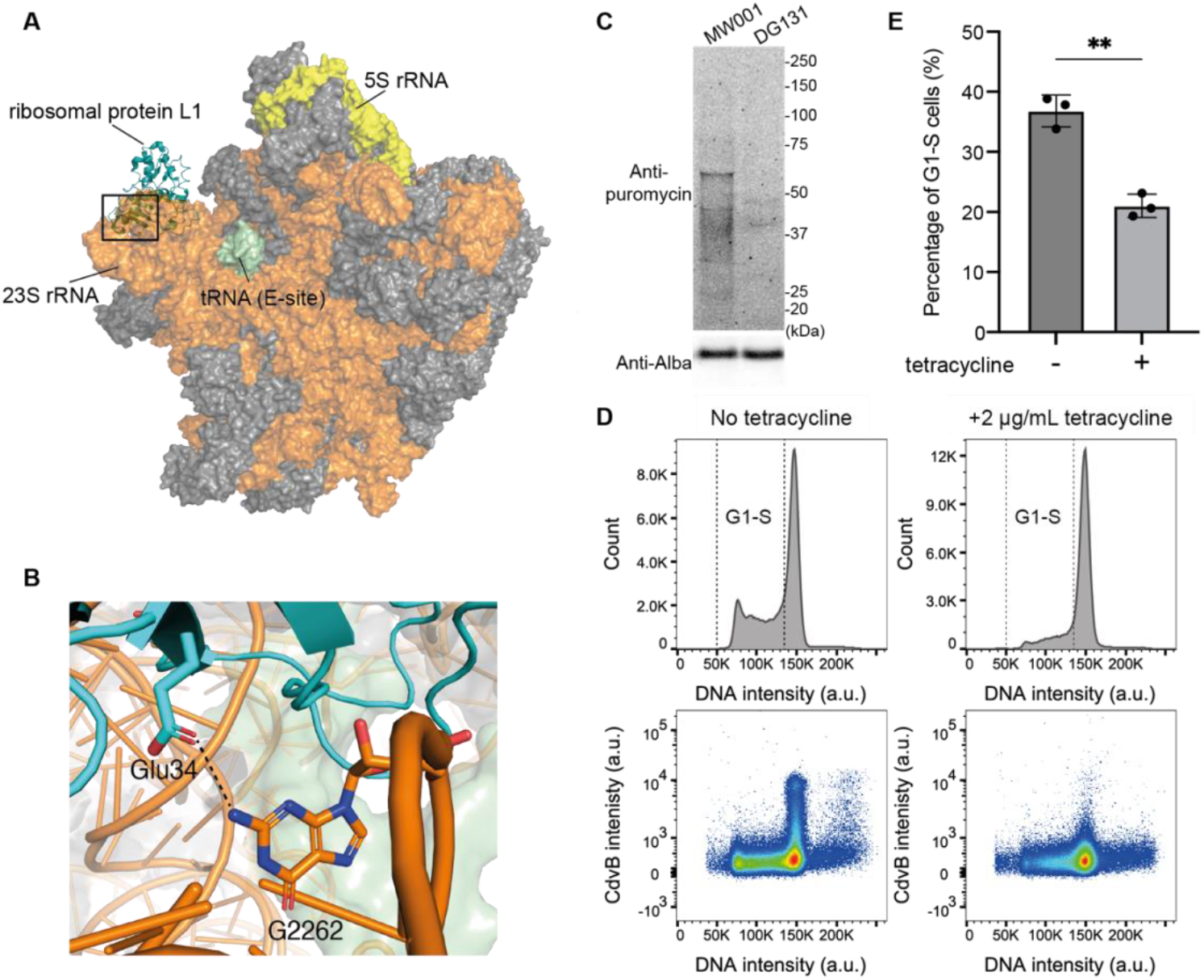
Translational inhibition results in G2 arrest in *S. acidocaldarius*. **(A)** Model showing the large ribosomal subunit Rpl1 (cyan), in relation to the 23S rRNA (orange) and 5S rRNA (yellow) subunits. Structure of the *S. acidocaldarius* 50S ribosome was obtained from PDB ID 8HKU (Wang *et al*., 2023). **(B)** The point mutation at E34K results in a reversal in charge, likely weakening the hydrogen bonding between Rpl1 and G2262 of 23S rRNA. Rpl1 sequence numbering is based on Uniprot P35024; 23S rRNA sequence numbering based on NCBI gene ID 3473376. This interface is highlighted in (A) in the box. **(C)** Puromycin incorporation is significantly reduced upon culture at the restrictive temperature (81 °C) in DG131. Shown is a representative Western blot with Alba as loading control. **(D)** Representative flow cytometry plots showing the DNA content and CdvB of control MW001 cells or MW001 cells treated with tetracycline. Inhibition of translation by tetracycline treatment results in a G2 arrest phenotype, similar to the TS phenotype of mutant DG131; n=2.5×10^5^ events. **(E)** Quantification of the percentage of G1-S cells in (D); no tetracycline: 36.8±2.6%, with tetracycline: 21.0±1.9% (mean ±SDs, N=3, Welch’s *t-*test *p=0*.*0016)*.

To test if the E34K mutation in Rpl1 disrupts ribosomal function at restrictive temperature to induce cell cycle arrest, we assayed translation activity in the DG131 strain by performing a puromycin incorporation assay at the permissive and restrictive temperature. This proved to be the case. Puromycin incorporation was severely restricted in DG131 when cultured at the restrictive temperature when compared to the uracil auxotrophic control strain MW001 (Fig. 3C). As a further test of this idea, we reasoned that if the TS phenotype of DG131 is due to translational inhibition, the treatment of control cells with the translational inhibitor, tetracycline (Cammarano *et al*., 1985; Aagaard *et al*., 1994), would induce a similar G2 arrest phenotype. Using the puromycin incorporation assay, we confirmed that tetracycline treatment strongly inhibited the translation activity in MW001 cells grown at 75 °C (Supplementary Fig. S1). Moreover, a 3-hour treatment with tetracycline significantly decreased the population of cells in G1-S phase of the cell cycle (Fig. 3D top panel, 3E) and reduced CdvB levels (Fig. 3D lower panel), phenocopying the effects of the E34K mutation at the restrictive temperature. Thus, translation inhibition induces a G2 arrest (Fig. 3D and 3E).

### Induction of wildtype Rpl1 expression in DG131 rescues its TS phenotype

In order to conclusively prove that the cell cycle arrest phenotype seen in DG131 cells at the restrictive temperature was due to the E34K point mutation in Rpl1, we attempted to rescue the phenotype by restoring the expression of wild-type Rpl1.

To ensure precise temperature control, we used a computer-controlled hotplate to enable automated temperature control and the gradual heating of samples (Fig. 4A). We then cultured strains in Duran bottles containing liquid media on a Hei-Connect hotplate with constant mixing using a magnetic stirrer. Ventilation was achieved using a 2.5mL serological pipette through a rubber stopper on the bottle cap, which allows the insertion of a probe enabling temperature-feedback control. In this experiment, DG131 cultures carrying either an empty vector or a vector containing a wild-type Rpl1 gene under the arabinose promoter were grown overnight at the permissive temperature to exponential phase. Protein expression was induced by the addition of arabinose for one hour. We then gradually increased the temperature to reach restrictive temperature after a period of 30 minutes. Finally, after another 7 hours at the restrictive temperature, cells were collected for fixation using ethanol and stained for the flow cytometry analysis (Fig. 4B). As before, DG131 cells expressing the empty vector underwent G2 arrest at the restrictive temperature as measured by DNA content (Fig. 4C and 4D). Strikingly, however, the expression of wild-type Rpl1 for an hour prior to temperature shift to the restrictive temperature was sufficient to prevent the arrest as measured by the restoration of the G1-S population (Fig. 4C and 4D).

**Figure 4.**
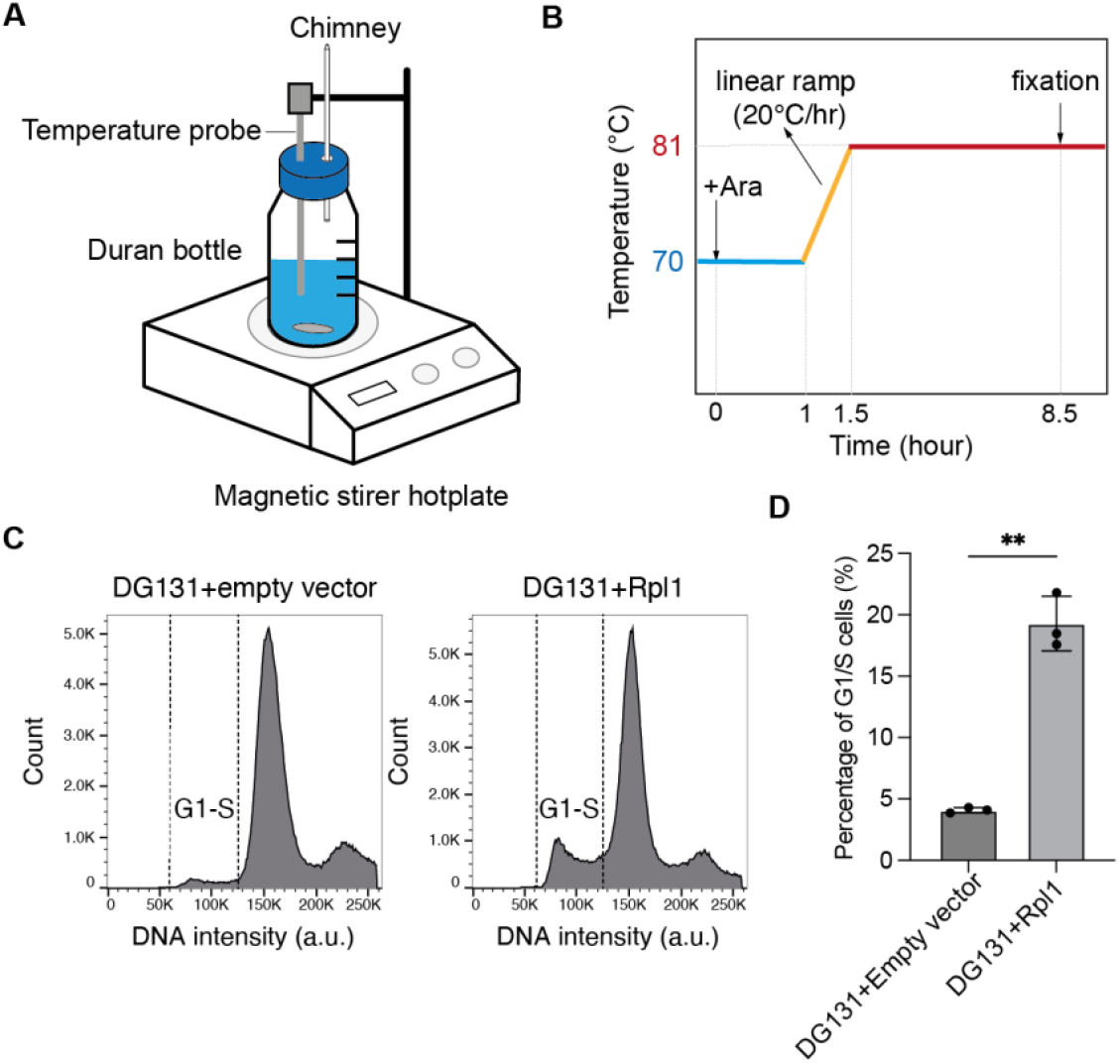
Rescue of DG131 TS phenotype. **(A)** Setup used for the accurate control of temperature in TS cultures. Strains were grown in Duran bottles on a magnetic stirrer hotplate with temperature control. The chimney serves as both a pathway for aeration and a condensation tube to reduce the evaporation of the media. **(B)** Plot showing the temperature shifts over the course of the experiment. Strains were cultured to the exponential phase of growth overnight at 70 °C before arabinose induction of protein expression for 1h. A linear ramp of 10 °C was initiated for 30min to the restrictive temperature of 81 °C, followed by 7 hours of culture. **(C)** Flow cytometry analysis of DNA content shows DG131 transformed with the empty vector displayed G2 arrest upon shift to the restrictive growth temperature, while DG131 expressing the wild-type Rpl1 retains its G1-S population. N=3, n=1×10^5^ events. **(D)** Quantification of the percentage of G1-S cells in (C); empty vector: 4.1±0.22%, Rpl1 rescue: 19.3±2.2% (mean ±SDs, N=3, Welch’s *t-*test *p=0*.*0066)*.

Taken together, these results show how forward genetic screens can be used as an approach to identify the genes involved in coupling cell growth and division in Sulfolobus, and demonstrate that translational inhibition arrests *S. acidocaldarius* in G2 of the cell cycle.

## Discussion

### Challenges of temperature sensitive mutant screen in Sulfolobus

Through this analysis, we have had to overcome several unexpected technical challenges that make a forward genetic analysis in *Sulfolobus* difficult.

First, for reproducibility when analysing mutant phenotypes in the presence and absence of a rescue construct, the rate of heating and the maintenance of a stable temperature proved to be crucial. While usage of a heating incubator allows simultaneous culturing of multiple strains during screening, the temperature variation inside the incubator can be large and the sampling process can also lead to transient changes in temperature that interfere with the experiment.

Second, the chemical mutagen must be carefully titrated. On the one hand, high mutation rates make it challenging to tease out the mutation(s) responsible for the observed phenotype using genomic sequencing. In the case of this screen, multiple mutations were detected in all mutant strains sequenced (Supplementary table S2). Note that in yeast screens sexual complementation can be used to simplify the problem by placing mutations into allelic groups to identify those responsible for a specific mutant phenotype (Hoffman *et al*., 2015). Conversely, the lower the rate of mutation, the larger the number of colonies that must be screened using replica plating to identify strains exhibiting a conditional mutant phenotype – something that is difficult to carry out at scale using *Sulfolobus*.

### Cell cycle phenotype of TS mutants

Further analysis of this set of TS mutants of *S. acidocaldarius* first isolated following chemical mutagenesis several decades ago by Dennis Grogan and Elaheh Mehdigholi has revealed several interesting observations. First, a majority of the TS strains displayed accumulation of G2 cells following a short incubation at the restrictive temperature, hinting that the transition from G2 into division phase (D-phase) is one of the most critical regulation points during cell cycle progression - functionally analogous to the G2/M checkpoint of eukaryotes (Stark and Taylor, 2004). This may also explain why whole genome sequencing of selected TS strains displaying G2 arrest phenotypes revealed mutations in genes with a wide range of functions, including metabolism and protein synthesis (Supplementary table S2). In agreement with the idea that defects in growth perturb the progression of cells from G2 of the cycle into division, we were able to show that the inhibition of translation activity via a temperature sensitive mutation of ribosome or by the treatment of cells with a translational inhibitor arrests cells in G2.

Similarly, previous studies have revealed that various conditions, such as acetic acid treatment, starvation, proteasome or topoisomerase inhibition (Bernander and Poplawski, 1997; Hjort and Bernander, 2001; Lundgren *et al*., 2004; Gadelle *et al*., 2005; Tarrason Risa *et al*., 2020) all prolong G2 or prevent cells from dividing. Thus, a wide range of stresses appears to induce an arrest that prevent cells from entering the division phase. While it is not clear why this is the case, this would seem to be a sensible place for cells with a 2N DNA content to arrest, given it enables cells to use the duplicate copies to repair DNA damage should it arise. In this light, it is interesting to note that like *Sulfolobus*, that haploid *S. pombe* cells have a short G1 phase, and tend to arrest in G2 following most insults, except when the arrest in G1 serves as a prelude to entry into meiosis (for example under conditions of nitrogen starvation) (Barnum and O’Connell, 2014; Yamashita *et al*., 2017; Vyas *et al*., 2021).

Interestingly, while there were also instances of TS mutants from the screen accumulating in S-phase cells at the restrictive temperature, we were unable to identify mutations that arrest cells in G1. This is in contrast to the situation in mammalian cells, where the G1/S transition is tightly regulated by cyclin/CDKs and E2F/Rb (Stark and Taylor, 2004; Barnum and O’Connell, 2014), causing cells to arrest at this stage of the cycle in response to a wide variety of stresses, including environmental factors and reductions in the rate of protein synthesis (Siede *et al*., 1994; Ivanova *et al*., 2011; Lockhead *et al*., 2020). Thus, a greater number of TS mutant lines will be needed to identify regulators of the G1/S transition.

Since many cell cycle synchronization experiments in *S. acidocaldarius* utilise acetic acid as an agent to arrest cells at G2 (Lundgren *et al*., 2004), which likely induces stress, our study suggests the possibility of using translational inhibition with tetracycline, or temperature shift of TS mutants (DG155 for example) as a useful alternative method for synchronizing cells to identify the consensus changes in multiomics levels during cell cycle progression. While this is the case, the mutation characterised in detail here proved irreversible, implying that cells accumulating defective ribosomes as the result of the Rpl1 mutation are unable to repair them following restoration of the permissive temperature.

In conclusion, in this study we show how a classical forward genetic screen can be used as a tool to identify genes involved in cell growth and division in archaea. Given the growing interest in the cell biology of archaea and their use as model organisms, the use of such phenotypic-driven screening of randomly induced mutations will be essential in expanding our understanding of archaea cell biology in the future.

## Materials and methods

### Strain and growth conditions

All cultures were grown in Brock medium supplemented with 0.1% (w/v) NZ-amine and 0.2% (w/v) sucrose (Brock *et al*., 1972), with 4μg/ml uracil supplemented as necessary. The permissive and restrictive temperatures used in this study are 70 °C and 81 °C respectively, unless otherwise stated.

All temperature sensitive (TS) *Sulfolobus acidocaldarius* mutants were derived from strain DG64 (*pyrB4*) (Grogan and Gunsalus, 1993; Bernander *et al*., 2000). TS strains were initially isolated by Bernander, Elaheh Mehdigholi, Grogan and colleagues as described previously (Bernander *et al*., 2000), and were cryogenically preserved for two decades prior to revival for this study. Strains were shipped from the University of Cincinnati, Ohio, USA, to the MRC Laboratory of Molecular Biology in Cambridge, UK, in sterilised 0.5mL microcentrifuge tubes at ambient temperature. Each microcentrifuge tube contains an agar plug cored from a xylose-tryptone-uracil plate and covered in confluent growth of a strain. Cores were dissolved by addition of warm Brock medium and transferred to 50mL falcon tubes containing 10mL of prewarmed Brock medium supplemented with uracil. Following three days of culture at 70 °C in a shaking incubator, cells were pelleted via centrifugation at 4000rcf for 5min, resuspended in Brock medium containing 50% glycerol (v/v), and frozen at -70 °C.

Frozen glycerol stocks of *S. acidocaldarius* were thawed by scraping with a sterile pipette tip and inoculating in 15mL of room temperature Brock medium. All cultures were passaged once, and then grown overnight to an OD_595_ of 0.10-0.25 at the start of all experiments. Protein expression in the rescue mutants were induced by addition of L-arabinose to the culture medium to a final concentration of 0.2% (w/v) in exponentially growing cultures.

### Temperature shift experiments

For the initial screen of cell cycle phenotypes in the TS mutant library, TS mutant strains were cultured in Erlenmeyer flasks at 70 °C overnight up to an OD_595_ of 0.10-0.25. This was split into two cultures, with one shifted to the restrictive temperature of 81 °C. After 6h of culture at the permissive and restrictive temperatures, cells were fixed by three stepwise additions of 4 °C ethanol to a final concentration of 70% and then stored at 4 °C, as described previously (Cezanne *et al*., 2023).

For the TS release experiments, TS mutants were cultured in Erlenmeyer flasks overnight at 70 °C up to an OD_595_ of 0.10-0.25, before shifting to the restrictive temperature of 81 °C for 4.5h. Next, the flasks were shifted back to the permissive temperature of 70 °C for a further 3h. Cells were sampled and fixed at intermediate timepoints as required, as described above.

DG131 rescue experiments were performed by culturing 150mL of cells in 250mL duran bottles on a Hei-Connect hotplate magnetic stirrer (Heidolph Scientific Products GmbH), see Figure 4A and 4B for schematic. Cells were grown at 70 °C with 250rpm stirring overnight to an OD_595_ of 0.10-0.25 prior to addition of arabinose for induction of protein expression. Following 1h of induction, the temperature was increased gradually over 30min to 81 °C, which was held for a further 7h. Cells were sampled and fixed at the start and end of the experiment, as described above.

### Molecular genetics

To generate rescue plasmids for the TS mutants, the pyrEF cassette of protein expression plasmid containing arabinose-inducible promoter, pSVAaraFX-Stop (van der Kolk *et al*., 2020), was replaced by pyrB gene from *Saccharolobus solfataricus*. Double digestion of pSVAara-FX-STOP plasmid with restriction enzymes XmaI and KpnI was performed, followed by gel extraction. *S. solfataricus pyrB* (SSO0614) with flanking sequences of atctaaccc and ggtaccgattat at the 5’- and 3’-ends respectively was custom synthesised as double stranded DNA fragment (gBlock from Integrated DNA Technology). The gBlock was digested by XmaI and KpnI, purified by PCR cleanup kit, and ligated with the digested vector backbone. This pSVAara-FX-STOP(pyrB) vector is then used for subsequent cloning of gene-of-interest.

DG131 rescue strains were generated as follows. Note that the annotated gene AAY80779.1 for the large ribosomal subunit protein Rpl1, available on the NCBI database is incorrect, as it displays an N terminal truncation (178aa). The full length Rpl1 protein is 221aa long (UniProt ID P35024). The Rpl1 open reading frame was amplified from wild type *S. acidocaldarius* strain DSM639 via PCR between the start and stop codons, using primer pair *aatccatggctGTGAAGAAAGTGTTAGCGGA* and *cgcctcgagCGCTCTTTTAACTTTTACAG*. The PCR fragment was flanked by NcoI and XhoI restriction sites, with two bases insertion (CT) following the NcoI site to ensure it remains in-frame with the internal start codon present on the NcoI site. This was cloned into a pSVAara-FX-STOP(pyrB) plasmid using standard molecular genetics methods, as described above.

### Electrocompetent cells

50mL of *S. acidocaldarius* cells were grown at 70 °C to an OD_595_ of 0.20-0.30. The culture was chilled on ice and cells pelleted via centrifugation at 2500rcf for 10min at 4 °C. The cells were washed thrice in 30mL of ice-cold 20mM sucrose and pelleted at 2500rcf for 10min at 4 °C. The cell pellet was next resuspended with ice-cold 20mM sucrose to a final OD_595_ of 10, split into 50μL aliquots and stored at -70 °C.

### Transformation of *S. acidocaldarius*

Transformation of *S. acidocaldarius* was performed following established protocols for *Sulfolobus* genetics (Wagner *et al*., 2012). Briefly, ER1821 *E. coli* was transformed with plasmid DNA for methylation in order to prevent restriction by the *SuaI* enzyme of *S. acidocaldarius* first. Methylated plasmids were next transformed into competent cells using a GeneGene Pulser Xcell (BioRad) at 2000V, 600Ω, 25μF in 1mm Gene Pulser Electroporation Cuvette (BioRad, 1652089). 1mL of Brock Medium without sucrose supplemented was then added and the cells incubated in a fresh microcentrifuge tube at 70 °C for 60min for recovery. Cells were then plated on two Brock medium plates containing 0.6% gelrite, without uracil supplemented, for 5 to 7 days at 70 °C. Single colonies were picked and grown up in liquid medium without uracil, and verified through genotyping. Positive clones were frozen in Brock Medium containing 50% glycerol (v/v) at -70 °C.

### Flow cytometry

1.5mL of fixed cells were collected via centrifugation at 8000rcf for 3min and the ethanol discarded. Cells were then washed twice in 1mL of phosphate-buffered saline containing 0.2% Tween-20 (Sigma-Aldrich, P9416) and 3% Bovine Serum Albumin (Sigma-Aldrich, A7030) (PBSTA). Cells were resuspended in primary antibodies (Supplementary table S3) in 200μL of PBSTA, and incubated overnight with shaking at room temperature on an Eppendorf Thermomixer. Next, cells were washed twice in 1mL of phosphate-buffered saline containing 0.2% Tween-20 (PBST) and resuspended in 200μL of PBSTA containing secondary antibodies (Supplementary table S3). This was incubated for 2h at room temperature with shaking on an Eppendorf Thermomixer. The cells were washed twice in 1mL PBST and then resuspended in 500μL of PBST containing 2μM Hoechst (Thermo Fisher Scientific, 62249) for labelling of DNA.

All flow cytometry experiments were performed on a BD Biosciences LSRFortessa. Laser wavelengths used are 355, 488, 561 and 640nm, in conjunction with the emission filters 450/50, 530/30, 586/15 and 670/14 respectively. All flow cytometry data was analysed in FlowJo v10 software.

### Whole genome sequencing

4 mL of cultures in stationary phase were collected by centrifugation and genomic DNA extraction was performed by phenol/chloroform extraction. Briefly, cell pellets were resuspended in 250μL of 10mM Tris-HCl pH8.0, 1mM EDTA, 150mM NaCl, 0.1% TritonX-100 (TENT) buffer in a microcentrifuge tube and incubated at room temperature for 10min. Next, 250μL of phenol:chloroform:isoamyl alcohol (25:24:1, v/v) (Invitrogen, 15593031) was added and the mixture vortexed for 30s. This was then centrifuged for 5min at 16,000rcf at room temperature. The upper aqueous phase was then carefully transferred to a fresh microcentrifuge tube and the phenol/chloroform extraction repeated an additional two times. 200μL of the final upper aqueous phase was then transferred to a fresh microcentrifuge tube, followed by addition of 150μL of 5M ammonium acetate and 875μL of 100% ethanol. The mixture was incubated overnight at -20 °C for DNA precipitation. Next, the precipitated DNA was pelleted by centrifugation at 16,000rcf for 5min, and the DNA pellet washed twice in ice-cold 70% ethanol. The ethanol was removed and the pellet air dried at room temperature for 10min. The DNA pellet was then resuspended in 50μL of nuclease free water and stored at 4 °C.

DNA concentration was determined using a Qubit dsDNA BR assay (Invitrogen, Q33265) on a Qubit Flex Fluorometer (Invitrogen), following the manufacturer’s instructions. Whole genome sequencing and mutation analysis was performed by Eurofins Genomics. In brief, the filtered sequence reads were mapped to the reference genome and subjected to variant analysis to identify mutations (SNP and InDel). Variant calling was performed by HalotypeCaller (Sentieon) with customised filters. The identified mutations were summarized in Table S2.

### Puromycin incorporation assay

The stock solution of puromycin (Sigma-Aldrich, P8833) was prepared in water with a concentration of 50mg/mL. MW001 and DG131 were cultured at the permissive temperature of 70 °C until OD_595_ of 0.10-0.25. The temperature was increased to the restrictive temperature of 81 °C for 3h, before the addition of 100μg/mL of puromycin for 15min. 5mL of cells were collected for Western blot analysis by centrifugation at 4000rcf for 5min at room temperature. The cell pellets were then washed by 1mL of room temperature BNS media followed by centrifugation of 20,000rcf for 1min. The supernatant was discarded and cell pellets frozen at -20 °C till further use.

### Tetracycline inhibition of translation

Stock solution of tetracycline was prepared in water with a concentration of 5mg/mL. Strains were treated with 2μg/mL tetracycline for 3h, before fixation for flow cytometry analysis as described above.

### Western blotting

Cell pellets were resuspended in 200μL of 1x NuPAGE LDS sample buffer (Invitrogen, NP0007) containing 10% β-mercaptoethanol and boiled for 5min. Next, the samples were centrifuged briefly and then loaded on a NuPAGE 4 to 12% Bis-Tris gels (Invitrogen, NP0321BOX). SDS-PAGE was performed at a constant 150V for 65min in MES running buffer. Proteins were next transferred to a nitrocellulose membrane at a constant 100V for 1h at 4 °C, before blocking in PBST containing 5% milk for 1h. Primary antibodies were added (Supplementary table S3) and blots were incubated overnight at 4 °C with gentle agitation. Following three washes in PBST for 5min on a rocking mixer, PBST containing 5% milk and IRDye-conjugated secondary antibodies (Supplementary Table S3) was added. This was allowed to incubate for 2h at room temperature, before three washes in PBST. Blots were imaged on a BioRad ChemiDoc system.

## Author contributions

All cell biology, genetics, biochemistry and flow cytometry experiments were performed by Y-WK and SF Revival of TS mutant stocks was performed by DWG. Initial screen of TS mutants was performed by JT, Y-WK and SF. Y-WK and SF co-wrote the manuscript. BB supervised the project and co-edited the manuscript with DWG.

## Acknowledgements

The authors would like to thank Joe Parham for his assistance in the culture and cryogenic preservation of TS mutant strains. Flow cytometry experiments were performed at the Medical Research Council Laboratory of Molecular Biology Flow Cytometry facility. We would like to thank members of the Flow Cytometry Facility for their assistance and technical support. Y-WK was supported by an EMBO postdoctoral fellowship (ALTF 903-2021) and by the Medical Research Council - Laboratory of Molecular Biology (MC_UP_1201/27); SF was supported by the Wellcome Trust (222460/Z/21/Z); BB received support for work in *Sulfolobus* from the Medical Research Council - Laboratory of Molecular Biology (MC_UP_1201/27), the Wellcome Trust (222460/Z/21/Z), and the Life Sciences Moore-Simons Foundation (735929LPI).

## References

Aagaard, C, Phan, H, Trevisanato, S, and Garrett, RA (1994). A spontaneous point mutation in the single 23S rRNA gene of the thermophilic arachaeon Sulfolobus acidocaldarius confers multiple drug resistance. J Bacteriol 176, 7744–7747.

Adames, NR, Gallegos, JE, and Peccoud, J (2019). Yeast genetic interaction screens in the age of CRISPR/Cas. Curr Genet 65, 307–327.

Albers, S-V, and Meyer, BH (2011). The archaeal cell envelope. Nat Rev Microbiol 9, 414–426.

Barnum, KJ, and O’Connell, MJ (2014). Cell Cycle Regulation by Checkpoints. Methods Mol Biol 1170, 29–40.

Bernander, R, and Poplawski, A (1997). Cell cycle characteristics of thermophilic archaea. J Bacteriol 179, 4963–4969.

Bernander, R, Poplawski, A, and Grogan, DW (2000). Altered patterns of cellular growth, morphology, replication and division in conditional-lethal mutants of the thermophilic archaeon Sulfolobus acidocaldarius. Microbiology 146, 749–757.

Boucher, Y, Kamekura, M, and Doolittle, WF (2004). Origins and evolution of isoprenoid lipid biosynthesis in archaea. Molecular Microbiology 52, 515–527.

Brock, TD, Brock, KM, Belly, RT, and Weiss, RL (1972). Sulfolobus: a new genus of sulfur-oxidizing bacteria living at low pH and high temperature. Arch Mikrobiol 84, 54–68.

Brügger, K, Redder, P, She, Q, Confalonieri, F, Zivanovic, Y, and Garrett, RA (2002). Mobile elements in archaeal genomes. FEMS Microbiol Lett 206, 131–141.

Cammarano, P, Teichner, A, Londei, P, Acca, M, Nicolaus, B, Sanz, JL, and Amils, R (1985). Insensitivity of archaebacterial ribosomes to protein synthesis inhibitors. Evolutionary implications. EMBO J 4, 811–816.

Cezanne, A, Foo, S, Kuo, Y-W, and Baum, B (2024). The Archaeal Cell Cycle. Annu Rev Cell Dev Biol.

Cezanne, A, Hoogenberg, B, and Baum, B (2023). Probing archaeal cell biology: exploring the use of dyes in the imaging of Sulfolobus cells. Front Microbiol 14, 1233032.

Charles-Orszag, A, Lord, SJ, and Mullins, RD (2021). High-Temperature Live-Cell Imaging of Cytokinesis, Cell Motility, and Cell-Cell Interactions in the Thermoacidophilic Crenarchaeon Sulfolobus acidocaldarius. Front Microbiol 12, 707124.

Chen, L, Brügger, K, Skovgaard, M, Redder, P, She, Q, Torarinsson, E, Greve, B, Awayez, M, Zibat, A, Klenk, H-P, et al. (2005). The genome of Sulfolobus acidocaldarius, a model organism of the Crenarchaeota. J Bacteriol 187, 4992–4999.

Gadelle, D, Bocs, C, Graille, M, and Forterre, P (2005). Inhibition of archaeal growth and DNA topoisomerase VI activities by the Hsp90 inhibitor radicicol. Nucleic Acids Res 33, 2310–2317.

Grabowski, B, and Kelman, Z (2001). Autophosphorylation of Archaeal Cdc6 Homologues Is Regulated by DNA. Journal of Bacteriology 183, 5459–5464.

Grogan, DW, Carver, GT, and Drake, JW (2001). Genetic fidelity under harsh conditions: analysis of spontaneous mutation in the thermoacidophilic archaeon Sulfolobus acidocaldarius. Proc Natl Acad Sci U S A 98, 7928–7933.

Grogan, DW, and Gunsalus, RP (1993). Sulfolobus acidocaldarius synthesizes UMP via a standard de novo pathway: results of biochemical-genetic study. J Bacteriol 175, 1500–1507.

Hjort, K, and Bernander, R (1999). Changes in cell size and DNA content in Sulfolobus cultures during dilution and temperature shift experiments. J Bacteriol 181, 5669–5675.

Hjort, K, and Bernander, R (2001). Cell cycle regulation in the hyperthermophilic crenarchaeon Sulfolobus acidocaldarius. Molecular Microbiology 40, 225–234.

Hoffman, CS, Wood, V, and Fantes, PA (2015). An Ancient Yeast for Young Geneticists: A Primer on the Schizosaccharomyces pombe Model System. Genetics 201, 403–423.

Ivanova, T, Gómez-Escoda, B, Hidalgo, E, and Ayté, J (2011). G1/S transcription and the DNA synthesis checkpoint: Common regulatory mechanisms. Cell Cycle 10, 912–915.

Iwabe, N, Kuma, K, Hasegawa, M, Osawa, S, and Miyata, T (1989). Evolutionary relationship of archaebacteria, eubacteria, and eukaryotes inferred from phylogenetic trees of duplicated genes. Proc Natl Acad Sci U S A 86, 9355–9359.

Iwai, T, Kurosawa, N, Itoh, YH, and Horiuchi, T (2000). Phylogenetic analysis of archaeal PCNA homologues. Extremophiles 4, 357–364.

van der Kolk, N, Wagner, A, Wagner, M, Waßmer, B, Siebers, B, and Albers, S-V (2020). Identification of XylR, the Activator of Arabinose/Xylose Inducible Regulon in Sulfolobus acidocaldarius and Its Application for Homologous Protein Expression. Front Microbiol 11, 1066.

Lake, JA, Henderson, E, Oakes, M, and Clark, MW (1984). Eocytes: a new ribosome structure indicates a kingdom with a close relationship to eukaryotes. Proc Natl Acad Sci U S A 81, 3786–3790.

Lindås, A-C, Chruszcz, M, Bernander, R, and Valegård, K (2014). Structure of crenactin, an archaeal actin homologue active at 90°C. Acta Cryst D 70, 492–500.

Liu, J, Gao, R, Li, C, Ni, J, Yang, Z, Zhang, Q, Chen, H, and Shen, Y (2017). Functional assignment of multiple ESCRT-III homologs in cell division and budding in Sulfolobus islandicus. Mol Microbiol 105, 540–553.

Lockhead, S, Moskaleva, A, Kamenz, J, Chen, Y, Kang, M, Reddy, AR, Santos, SDM, and Ferrell, JE (2020). The Apparent Requirement for Protein Synthesis during G2 Phase Is due to Checkpoint Activation. Cell Reports 32.

Lundgren, M, Andersson, A, Chen, L, Nilsson, P, and Bernander, R (2004). Three replication origins in Sulfolobus species: synchronous initiation of chromosome replication and asynchronous termination. Proc Natl Acad Sci U S A 101, 7046– 7051.

Meyer, BH, Adam, PS, Wagstaff, BA, Kolyfetis, GE, Probst, AJ, Albers, SV, and Dorfmueller, HC (2022). Agl24 is an ancient archaeal homolog of the eukaryotic N-glycan chitobiose synthesis enzymes. eLife 11, e67448.

Nikulin, A, Eliseikina, I, Tishchenko, S, Nevskaya, N, Davydova, N, Platonova, O, Piendl, W, Selmer, M, Liljas, A, Drygin, D, et al. (2003). Structure of the L1 protuberance in the ribosome. Nat Struct Biol 10, 104–108.

Pulschen, AA, Mutavchiev, DR, Culley, S, Sebastian, KN, Roubinet, J, Roubinet, M, Risa, GT, Wolferen, M van Roubinet, C, Schmidt, U, et al. (2020). Live Imaging of a Hyperthermophilic Archaeon Reveals Distinct Roles for Two ESCRT-III Homologs in Ensuring a Robust and Symmetric Division. Current Biology 30, 2852-2859.e4.

She, Q, Singh, RK, Confalonieri, F, Zivanovic, Y, Allard, G, Awayez, MJ, Chan-Weiher, CC, Clausen, IG, Curtis, BA, De Moors, A, et al. (2001). The complete genome of the crenarchaeon Sulfolobus solfataricus P2. Proc Natl Acad Sci U S A 98, 7835–7840.

Siede, W, Friedberg, AS, Dianova, I, and Friedberg, EC (1994). Characterization of G1 checkpoint control in the yeast Saccharomyces cerevisiae following exposure to DNA-damaging agents. Genetics 138, 271–281.

Stark, GR, and Taylor, WR (2004). Analyzing the G2/M checkpoint. Methods Mol Biol 280, 51–82.

Tarrason Risa, G. Hurtig, F, Bray, S, Hafner, AE, Harker-Kirschneck, L, Faull, P, Davis, C, Papatziamou, D, Mutavchiev, DR, Fan, C, et al. (2020). The proteasome controls ESCRT-III-mediated cell division in an archaeon. Science 369, eaaz2532.

Vyas, A, Freitas, AV, Ralston, ZA, and Tang, Z (2021). Fission Yeast Schizosaccharomyces pombe: A Unicellular “Micromammal” Model Organism. Curr Protoc 1, e151.

Wagner, M, van Wolferen, M, Wagner, A, Lassak, K, Meyer, BH, Reimann, J, and Albers, S-V (2012). Versatile Genetic Tool Box for the Crenarchaeote Sulfolobus acidocaldarius. Front Microbiol 3, 214.

Wang, Y-H, Dai, H, Zhang, L, Wu, Y, Wang, J, Wang, C, Xu, C-H, Hou, H, Yang, B, Zhu, Y, et al. (2023). Cryo-electron microscopy structure and translocation mechanism of the crenarchaeal ribosome. Nucleic Acids Res 51, 8909–8924.

Woese, CR, and Fox, GE (1977). Phylogenetic structure of the prokaryotic domain: the primary kingdoms. Proc Natl Acad Sci U S A 74, 5088–5090.

Yamashita, A, Sakuno, T, Watanabe, Y, and Yamamoto, M (2017). Analysis of Schizosaccharomyces pombe Meiosis. Cold Spring Harb Protoc 2017, pdb.top079855.

Zhang, C, Phillips, APR, Wipfler, RL, Olsen, GJ, and Whitaker, RJ (2018). The essential genome of the crenarchaeal model Sulfolobus islandicus. Nat Commun 9, 4908.

Zheng, J, Mallon, J, Lammers, A, Rados, T, Litschel, T, Moody, ERR, Ramirez-Diaz, DA, Schmid, A, Williams, TA, Bisson-Filho, AW, et al. (2023). Salactin, a dynamically unstable actin homolog in Haloarchaea. mBio 0, e02272–23.

Zink, IA, Pfeifer, K, Wimmer, E, Sleytr, UB, Schuster, B, and Schleper, C (2019). CRISPR-mediated gene silencing reveals involvement of the archaeal S-layer in cell division and virus infection. Nat Commun 10, 4797.

