## Supplemental materials for "A temperature sensitive mutant screen reveals translational stress-induced cell cycle regulation in a thermophilic archaeon"

**Supplementary Table S1: Summary of TS mutants**

| Mutant Class | Description | Occurrence | Strain Number |
|---|---|---|---|
| Class 1 | Mild phenotype | 22/44 | DG113, DG114, DG119, DG121, DG123, DG124, DG125, DG126, DG127, DG128, DG130, DG133, DG134, DG138, DG141, DG142, DG143, DG144, DG147, DG149, DG152, DG153 |
| Class 2 | G2 arrest | 11/44 | DG115, DG116*, DG129, DG131*, DG132, DG137, DG139, DG154*, DG155*, DG156, DG157 |
| Class 3 | S-phase phenotype | 3/44 | DG118, DG120*, DG140* |
| Class 4 | Defects at 70°C | 2/44 | DG135, DG151* |
| Conditional lethals | Lethal at 81°C | 6/44 | DG117, DG122, DG145, DG146, DG148, DG150 |

*See supplementary table S2 for sequencing results

**Supplementary Table S2. Summary of mutations in selected temperature sensitive mutants.** Purified genomic DNA was sequenced by next generation sequencing and the filtered sequencing reads were aligned to the reference genome (NCBI database GCF_002215565.1) to identify mutations. Mutations that were found in multiple strains are highlighted in gray. The point mutation of Rpl1 in DG131 is highlighted in yellow.

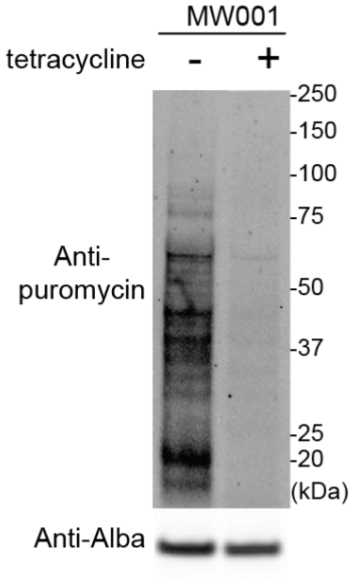


**Supplementary Figure S1. Tetracycline inhibits protein translation in *S. acidocaldarius*.** A non-temperature sensitive control strain of *S. acidocaldarius* (MW001) was treated with 2 µg/mL (right lane) or no tetracycline (left lane) for 3 hours at 75 °C, followed by treatment of 100 µg/mL of puromycin for 15 min. The incorporation of puromycin assayed by western blot was strongly reduced in the tetracycline-treated cells, suggesting that tetracycline acts as a translational inhibitor.

**Supplementary Table S3: Antibodies used in this study**

| Antibody | Host organism | Dilution | Catalog number |
|---|---|---|---|
| Anti-CdvB serum | Rabbit | 1:1000 | - |
| Anti-CdvB1 IgY | Chicken | 1:1000 | - |
| Anti-Puromycin IgG | Mouse | 1:2500 | ZMS1016 (Sigma-Aldrich) |
| Anti-Alba serum | Rabbit | 1:2000 | - |
| Anti-Rabbit IgG, AF488 | Goat | 1:1000 | A11034 (Invitrogen) |
| Anti-Chicken IgY, AF546 | Goat | 1:1000 | A11040 (Invitrogen) |
| Anti-Mouse IgG IRDye 800CW | Goat | 1:10,000 | 926-32210 (LI-COR Biosciences) |
| Anti-Rabbit IgG IRDye 800CW | Goat | 1:10,000 | 926-32211 (LI-COR Biosciences) |
